# Group IIA secreted phospholipase A_2_ in human serum kills commensal but not clinical *Enterococcus faecium* isolates

**DOI:** 10.1101/272617

**Authors:** Fernanda L. Paganelli, Helen L. Leavis, Samantha He, Nina M. van Sorge, Christine Payré, Gérard Lambeau, Rob J.L. Willems, Suzan H.M. Rooijakkers

## Abstract

Human innate immunity employs cellular and humoral mechanisms to facilitate rapid killing of invading bacteria. The direct killing of bacteria by human serum is mainly attributed to the activity of the complement system that forms pores in Gram-negative bacteria. Although Gram-positive bacteria are considered resistant to serum killing, we here uncover that normal human serum effectively kills *Enterococcus faecium.* Comparison of a well-characterized collection of commensal and clinical *E. faecium* isolates revealed that human serum specifically kills commensal *E. faecium* strains isolated from normal gut microbiota, but not clinical isolates. Inhibitor studies show that the human group IIA secreted phospholipase A2 (hGIIA), but not complement, is responsible for killing of commensal *E. faecium* strains in human normal serum. This is remarkable since hGIIA concentrations in ‘non-inflamed’ serum were considered too low to be bactericidal against Gram-positive bacteria. Mechanistic studies showed that serum hGIIA specifically causes permeabilization of commensal *E. faecium* membranes. Altogether, we find that a normal serum concentration of hGIIA effectively kills commensal *E. faecium* and that hGIIA resistance of clinical *E. faecium* could have contributed to the ability of these strains to become opportunistic pathogens in hospitalized patients.

**Importance:** Human normal serum contains antimicrobial components that effective kill invading Gram-negative bacteria. Although Gram-positive bacteria are generally considered resistant to serum killing, here we show that normal human effectively kills the Gram-positive *Enterococcus faecium* strains that live as commensals in the gut of humans. In contrast, clinical *E. faecium* strains that are responsible for opportunistic infections in debilitated patients are resistant against human serum. The key factor in serum responsible for killing is group IIA secreted phospholipase A2 (hGIIA) that effectively destabilizes commensal *E. faecium* membranes. We believe that hGIIA resistance by clinical *E. faecium* could have contributed to the ability of these strains to cause opportunistic infections in hospitalized patients. Altogether, understanding mechanisms of immune defense and bacterial resistance could aid in further development of novel anti-infective strategies against medically important multidrug resistant Gram-positive pathogens.

## Introduction

The human immune system is essential to protect us against invading bacterial infections. The first line of immune defense is comprised of cellular and humoral factors that together fight infections in the first minutes to hours of an infection against bacteria. Phagocytic cells, such as neutrophils are able to engulf invading bacteria by bacterial recognition, which is enhanced by bacterial labeling with serum factors as antibodies and complement activation products (1). However, human serum also harbors various antimicrobial proteins and peptides that can directly lyse bacteria without the help of immune cells (2). This serum bactericidal activity is mainly effective against Gram-negative bacteria that are sensitive for the pore-forming Membrane Attack Complex (MAC) of the human complement system (3). For long it has been known that Gram-positive bacteria are resistant to this ‘serum bactericidal activity’. It is generally assumed that all Gram-positive bacteria (both pathogenic and non-pathogenic) protect themselves against complement-induced pore formation via a thick layer of peptidoglycan that surrounds the bacterial membrane (4). We here found that normal human serum selectively kills commensal *E. faecium* strains, whereas disease-associated *E. faecium* strains remain viable.

*Enterococcus faecium* is a common inhabitant of the gut of mammals, birds and insects (5, 6). While *E. faecium* colonization is harmless in healthy individuals, this bacterium can cause serious infections in immunocompromised patients, such as bacteremia and endocarditis (7). In fact, multidrug resistant *E. faecium*, most notably vancomycin-resistant *E. faecium* (VRE), has emerged as an important cause of nosocomial infections worldwide (8). Recent work showed that clinical *E. faecium* isolates are genetically distinct from commensal strains (9). Whole genome comparison and functional assays identified various virulence factors and carbohydrate metabolism gene clusters that were enriched in clinical isolates and that could contribute to successful niche adaptation in hospitals (9–12). Hospitalized patients suffering from clinical *E. faecium* infections, frequently have a severely compromised cellular immunity. Yet, innate humoral immunity could still aid in *E. faecium* infection control (13). To what extent this is relevant in this infection type (by clinical *E. faecium* strains), is currently not completely understood (14–19).

Screening of a collection of commensal and multidrug resistant clinical *E. faecium* strains revealed that normal human serum specifically kills commensal strains. We found that the group IIA secreted phospholipase A2 (hGIIA) is key to the bactericidal activity of serum and that resistance to serum hGIIA could have contributed to the development of clinical *E. faecium* isolates.

## Results

### Human serum kills commensal but not clinical *Enterococcus faecium* strains

While human serum effectively kills Gram-negative bacteria (20), it is generally accepted that serum does not kill pathogenic and non-pathogenic Gram-positives (21). Serendipitously however, we discovered that a commensal strain of *E. faecium* (E1007, isolated from feces of a healthy individual) was effectively killed by human serum. We incubated 10^5^ exponential phase *E. faecium* E1007 with normal human serum (pooled from 20 healthy volunteers) and quantified bacterial survival via colony enumeration on agar plates (Fig 1A). Within 30 minutes, 10% human serum could completely kill 10^5^ E1007 bacteria. In contrast, a clinical *E. faecium* strain (E1162, isolated from blood of a hospitalized patient) was fully resistant to serum killing in a similar assay. Based on these results, we extended these analyses to a broader panel of genomically well-characterized clinical and commensal *E. faecium* isolates. We selected clinical *E. faecium* strains isolated from hospitalized patients and compared their serum susceptibility to a selection of commensal *E. faecium* isolated from healthy individuals or animals. Exposure of all 19 strains (described in Table 1) to 25% human serum revealed that human serum specifically kills commensal strains by 70% reduction on average, but not clinical *E. faecium* strains, which survive for 98% (Fig 1B).

**Table 1.**
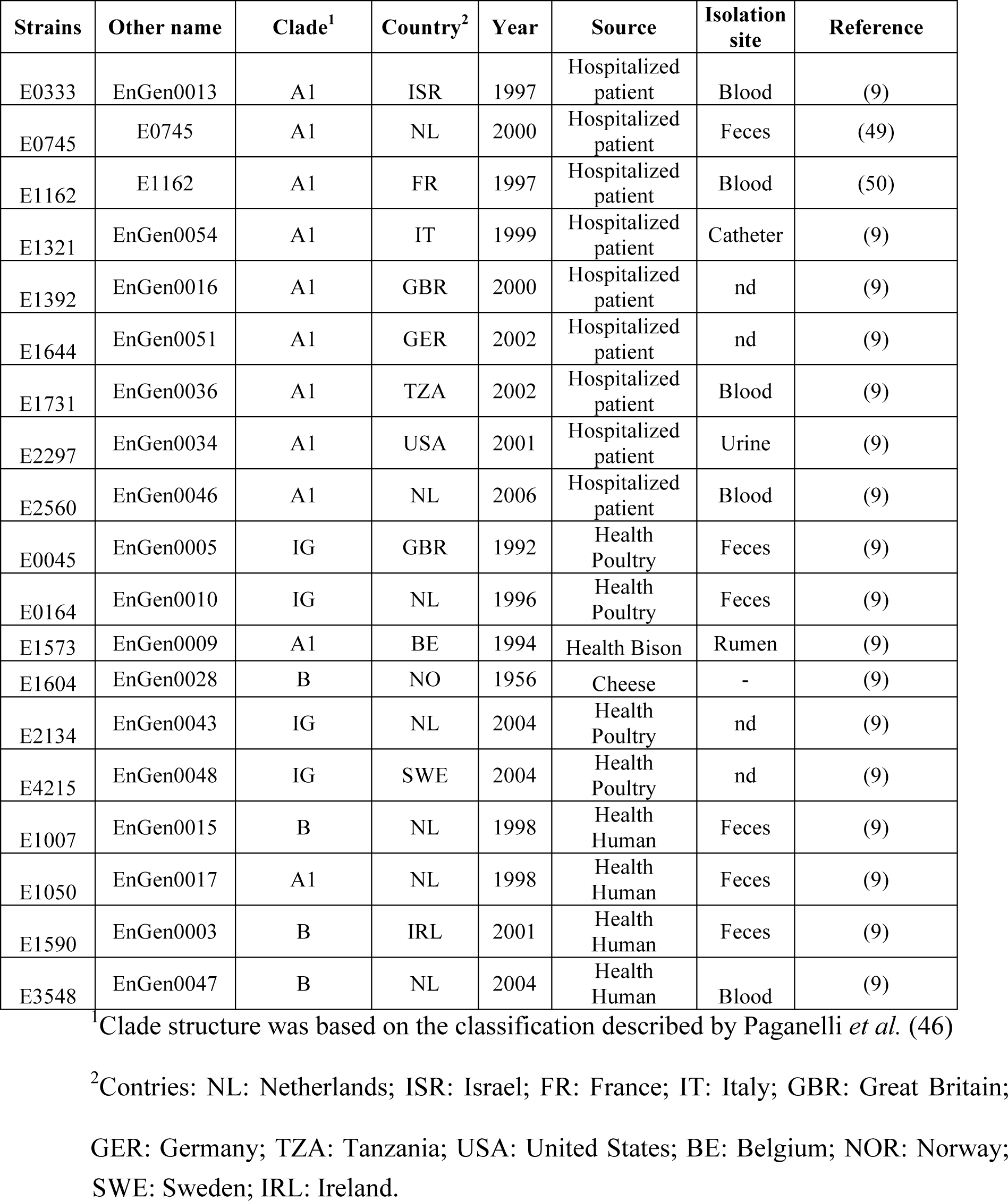
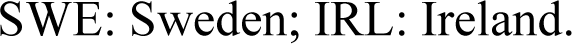
Ecological origin and clade assignment of 19 *E. faecium* strains.

| Strains | Other name | Clade<sup>1</sup> | Country<sup>2</sup> | Year | Source | Isolation site | Reference |
| --- | --- | --- | --- | --- | --- | --- | --- |
| E0333 | EnGen0013 | A1 | ISR | 1997 | Hospitalized patient | Blood | (9) |
| E0745 | E0745 | A1 | NL | 2000 | Hospitalized patient | Feces | (49) |
| E1162 | E1162 | A1 | FR | 1997 | Hospitalized patient | Blood | (50) |
| E1321 | EnGen0054 | A1 | IT | 1999 | Hospitalized patient | Catheter | (9) |
| E1392 | EnGen0016 | A1 | GBR | 2000 | Hospitalized patient | nd | (9) |
| E1644 | EnGen0051 | A1 | GER | 2002 | Hospitalized patient | nd | (9) |
| E1731 | EnGen0036 | A1 | TZA | 2002 | Hospitalized patient | Blood | (9) |
| E2297 | EnGen0034 | A1 | USA | 2001 | Hospitalized patient | Urine | (9) |
| E2560 | EnGen0046 | A1 | NL | 2006 | Hospitalized patient | Blood | (9) |
| E0045 | EnGen0005 | IG | GBR | 1992 | Health Poultry | Feces | (9) |
| E0164 | EnGen0010 | IG | NL | 1996 | Health Poultry | Feces | (9) |
| E1573 | EnGen0009 | A1 | BE | 1994 | Health Bison | Rumen | (9) |
| E1604 | EnGen0028 | B | NO | 1956 | Cheese | - | (9) |
| E2134 | EnGen0043 | IG | NL | 2004 | Health Poultry | nd | (9) |
| E4215 | EnGen0048 | IG | SWE | 2004 | Health Poultry | nd | (9) |
| E1007 | EnGen0015 | B | NL | 1998 | Health Human | Feces | (9) |
| E1050 | EnGen0017 | A1 | NL | 1998 | Health Human | Feces | (9) |
| E1590 | EnGen0003 | B | IRL | 2001 | Health Human | Feces | (9) |
| E3548 | EnGen0047 | B | NL | 2004 | Health Human | Blood | (9) |

<sup>1</sup>Clade structure was based on the classification described by Paganelli *et al.* (46)
<sup>2</sup>Contries: NL: Netherlands; ISR: Israel; FR: France; IT: Italy; GBR: Great Britain;
GER: Germany; TZA: Tanzania; USA: United States; BE: Belgium; NOR: Norway;
SWE: Sweden; IRL: Ireland.

**Figure 1.**
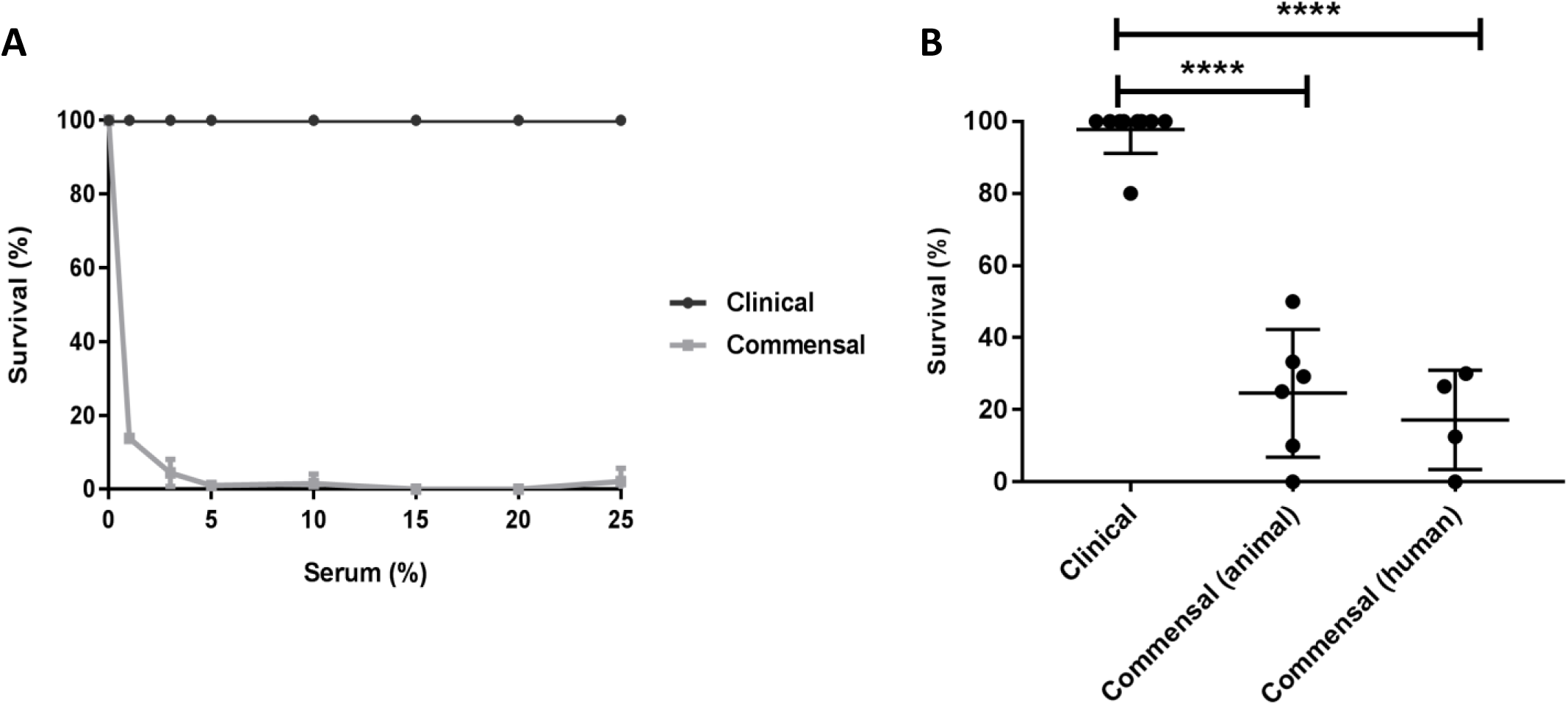
Human serum kills commensal, but not clinical *E. faecium* isolates. (A) Comparison of bacterial survival at different concentrations of pooled normal human serum of clinical *E. faecium* strain E1162 (clinical) and commensal *E. faecium* strain E1007 (commensal). (B) Killing of *E. faecium* isolates originating from hospitalized patients (clinical), healthy animals (commensal animal) and healthy humans (commensal humans) in 25% pooled human serum. Killing was quantified by comparison of total colony forming units after 30 min incubation in the buffer negative control (RPMI) compared to 25% human serum. Epidemiological details of *E. faecium* strains are listed in table 1.

### Complement does not contribute to serum-mediated killing of commensal *E. faecium*

To investigate how human serum kills commensal *E. faecium* strains, we first used several approaches to inactivate the complement system. Bactericidal activity of the complement system is mediated by the Membrane Attack Complex (MAC), a ring-structured pores consisting of proteins C5b, C6, C7, C8 and multiple copies of C9 (C5b-9). Formation of pores in bacterial membranes occurs rapidly following an enzymatic chain reaction on the cell surface. Initial experiments seemed to suggest that complement indeed played a role in killing of *E. faecium*. For instance, we found that *E. faecium* killing could be blocked by exposing the serum to 56°C (heat-inactivated (HI) serum), a method commonly used to inactivate certain heat-labile complement components (22) (Fig 2A-B). Furthermore, addition of the chelating agent (Ethylenediaminetetraacetic acid - EDTA), known to block the complement reaction (23), also interfered with serum killing (Fig 2A, B). However, when we used more specific inhibitors to block the complement reaction, we found that these inhibitors did not affect serum-mediated killing of *E. faecium* (Fig 2A). For instance, application of C3 inhibitor CP40 (24) and C5 inhibitor OmCI (25) did not block bactericidal activity (Fig 2A), although both inhibitors effectively blocked MAC deposition on the bacterial surface (Fig 2B). Resistance of the clinical *E. faecium* strain E1162 was not affected by any of the serum treatments (S1 Fig). From these data, we concluded that killing of commensal *E. faecium* in human serum is not mediated by complement, but by another heat-sensitive and divalent cation-dependent factor (26).

**Figure 2.**
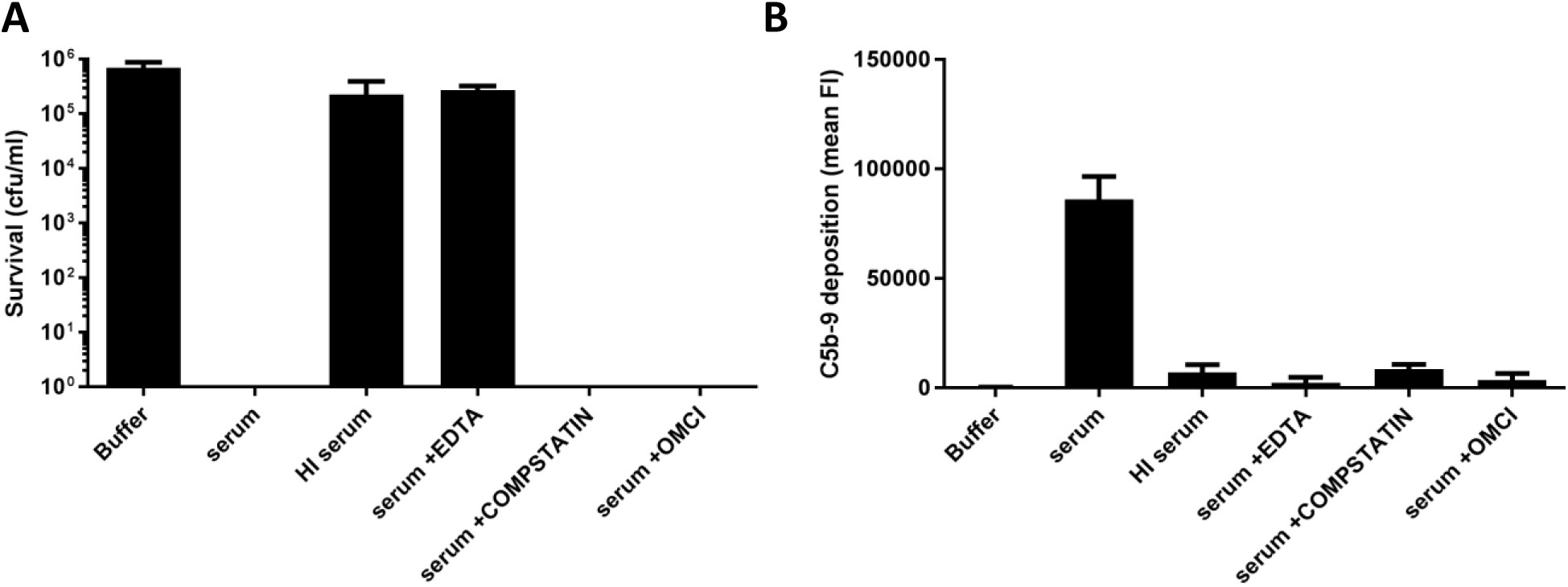
Complement does not contribute to serum-mediated killing of commensal *E. faecium.* (A) Killing of commensal *E. faecium* strain E1007 was measured by bacterial viability on blood agar plates after 30 min incubation at the indicated conditions. (B) Complement system activity in human serum was measured by C5b-9 deposition on bacterial surface of clinical isolate E1162.

### hGIIA is essential for serum killing of commensal *E. faecium*

Next, we decided to test the role of hGIIA (14 kDa), which catalytic function is calcium-dependent (27). hGIIA belongs to the secreted PLA2 family of enzymes present in various genomes, from humans to snakes, invertebrates, plants, fungi and even bacteria that hydrolyze membrane phospholipids at the *sn-2* position (28). Although immune cell derived hGIIA was previously identified as a bactericidal component against several Gram-positive bacteria(29), hGIIA concentrations in normal serum are much lower than the reported bactericidal concentrations for most Gram-positive bacteria (30–33). In our pooled human serum, we observed a concentration of hGIIA of 5 ng/ml (0.3 nM), similar to those described previously (31, 34) (S2 Fig). Nevertheless, we tested whether a specific hGIIA inhibitor (LY311727, Sigma-Aldrich) could block killing of commensal *E. faecium* in human serum. We observed that the inhibitor blocks the killing activity in a dose dependent manner (Fig 3A) and at different concentrations of human serum (Fig 3B). Moreover, the hGIIA inhibitor blocked killing of all tested commensal *E. faecium* strains (Fig 3C). Complementary, we could restore killing of commensal *E. faecium* in heat-inactivated serum by reconstituting serum with pure recombinant human hGIIA (Fig 3D). In summary, we found that hGIIA is the key player in killing of commensal *E. faecium* strains by normal human serum.

**Figure 3.**
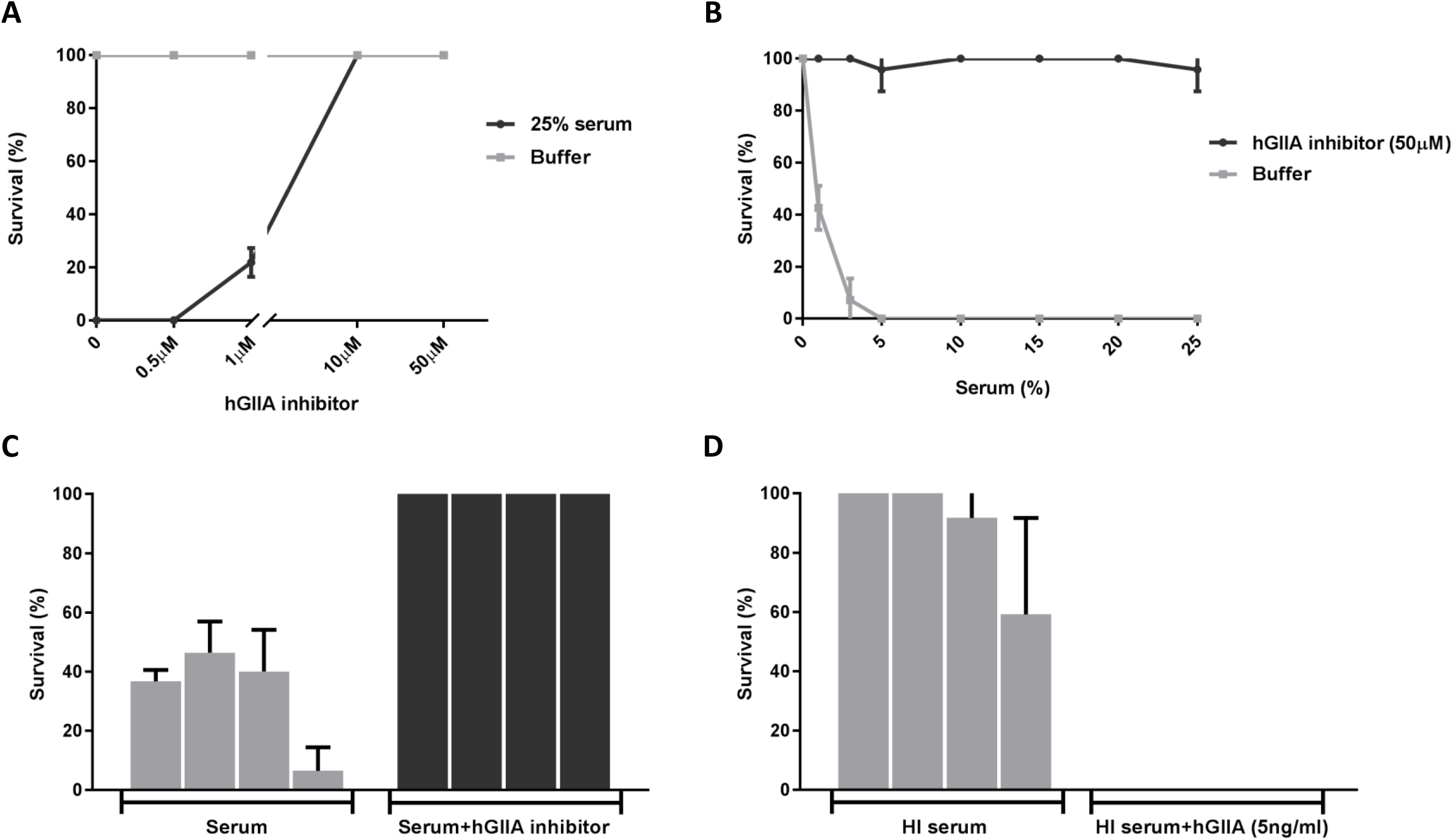
hGIIA is essential for serum killing of commensal *E. faecium.* (A) Killing of commensal *E. faecium* strain E1007 in 25% pooled human serum is inhibited by hGIIA inhibitor (LY311727) in a dose-dependent manner. (B) Killing of commensal *E. faecium* strain E1007 in different concentrations of human serum incubated with or without hGIIA inhibitor (LY311727). (C) Killing of *E. faecium* commensal strains E1050, 1590, E3548 and E1007 in 25% pooled human serum with or without 50 µM of hGIIA inhibitor (LY311727), respectively. (D) Killing of *E. faecium* commensal strains E1050, 1590, E3548 and E1007 in 25% pooled heat-inactivated (HI) human serum without or with recombinant hGIIA, respectively. Killing was quantified by bacterial viability on blood agar plates after 30 min incubation in the different conditions.

### Serum hGIIA causes membrane destabilization in commensal *E. faecium*

Finally, we studied how hGIIA in serum causes bacterial killing. hGIIA acts by hydrolyzing membrane phospholipids in bacterial membranes, whereas host cells are highly resistant to its activity at normal physiological concentrations (30). Here, we used a flow cytometric approach to study whether hGIIA can induce membrane damage to commensal *E. faecium* strains. Bacteria were incubated with human serum in the presence of the DNA dye (sytox green) that only binds DNA and RNA when bacterial membranes are damaged (35). While untreated *E. faecium* cells remain sytox negative, 90% in average of the commensal *E. faecium* population became sytox positive after 30 min incubation with serum concentrations equal or above 10% (Fig 4B). No increase in sytox intensity was observed in the clinical strain E1162 (Fig 4A-B). Flow cytometric quantification of sytox influx showed a concentration-dependent effect of serum at inducing membrane damage of commensal, but not clinical *E. faecium* strains (Fig 4B). Confocal microscopy confirmed membrane damage of the commensal *E. faecium* E1007 strain but not clinical E1162 strain by human serum (Fig 4C) with different DNA dyes (syto9 and propidium iodide). Furthermore, we found that the serum-induced membrane permeabilization of the E1007 strain was mediated by hGIIA, since EDTA and hGIIA inhibition blocked the observed sytox influx (Fig 4D). Finally, we found that all tested commensal, but not clinical *E. faecium* isolates are sensitive to membrane permeabilization via hGIIA in human serum (Fig 4E).

**Figure 4.**
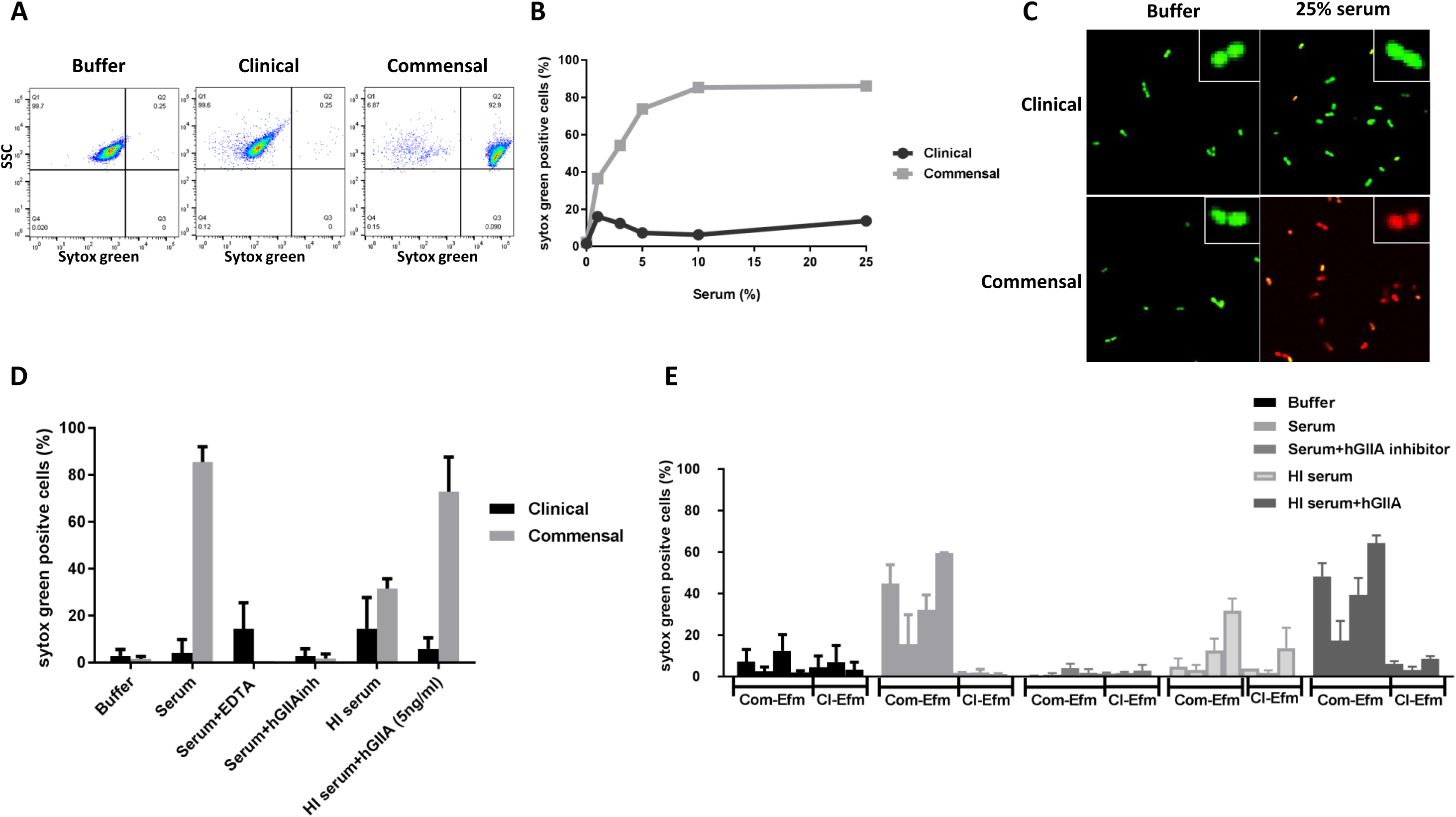
Serum hGIIA causes membrane permeabilization in commensal *E. faecium*. Representative flow cytometry plots of sytox green influx (switch to the right) of commensal *E. faecium* isolate E1007 incubated with RPMI (buffer) and clinical (E1162) and commensal (E1007) strains incubated with 25% pooled human serum. Sytox green influx in E1007 and E1162 using different concentration of human serum. (C) Confocal pictures of clinical (E1162) and commensal (E1007) *E. faecium* strains incubated with RPMI (buffer) or 25% human serum in the presence of syto9 (in green) representing live cells and propidium iodide (in red) representing damaged bacterial membrane. *E. faecium* membrane permeabilization of clinical strain E1162 and commensal strain E1007 was measured by sytox green influx in the described conditions. (D) Sytox green influx upon incubation in serum in the absence or presence of hGIIA inhibitors and upon restoration in heat-inactivated (HI) human serum with recombinant hGIIA. (E) Sytox green influx in *E. faecium* clinical strains E2560, E0745, E1162 and commensal strains E1050, E1590, E3548, E1007 in RPMI (buffer), 25% pooled human serum without or with 50 µM of hGIIA inhibitor (LY311727) or heat-inactivated (HI) human serum with 5 ng/ml recombinant hGIIA, respectively.

Altogether we conclude that hGIIA is the principal component of human serum, which effectively kills commensal *E. faecium* at low concentrations (in the subnanomolar nM range) by destabilization of the bacterial membrane.

## Discussion

Novel anti-infective strategies are pivotal to curtail the emergence of multidrug-resistant pathogens. Besides development of direct antibacterial compounds, drugs acting at the level of interaction between microbes and the immune system could be promising. Previous studies revealed that *E. faecium* can adopt distinct lifestyles (commensal and pathogenic), which are represented by two distinct genomic clades that differ in genetic polymorphisms and gene repertoire (9, 10). Here we identify that these two clades display distinct susceptibility to killing by normal human serum. Given the genetic variances in genes that are conserved in commensal and clinical strains as well as the differences in gene repertoire between clinical and commensal *E. faecium* strains, it is likely that gene expression and/or differences in gene content (a result of gene gain and loss) explain this difference in hGIIA resistance between commensal and clinical strains. Further investigation is needed to identify the bacterial factors that contribute to *E. faecium* resistance or susceptibility to serum, but genes related to changes in bacterial surface charge and involved in lipid synthesis in bacteria (36, 37) would be good candidates to be investigated, since hGIIA binding is directly influence by the negative charge of the bacterial surface and phospholipids modifications (30). We hypothesize that the development of human serum resistance may have contributed to the decisive step for *E. faecium* to evolve into a relevant pathogen. On the other hand, the ability of hGIIA to kill commensal Gram-positives can be an evolutionary strategy of the innate immune system to contain commensal bacteria in their specific niches and preclude invasion of sterile tissues.

Our study also highlights an important role for hGIIA in the humoral immune response against *E. faecium*. While other groups reported that high concentrations of hGIIA (produced locally in lungs and tears (38, 39) or in serum under septic shock conditions) (40, 41) could disrupt certain Gram-positive bacteria, its importance in normal serum or plasma has not been recognized in physiological concentrations, besides against *Listeria monocytogenes* (34). We observed restoration of killing of commensal *E. faecium* when recombinant hGIIA was added in physiological concentrations in heat-inactivated human serum. Although our data indicate that hGIIA is the major factor in serum responsible for the killing phenotype observed against commensal *E. faecium* strains, further studies are necessary to identify the specific conditions in serum that facilitate hGIIA action.

hGIIA is known for its ability to hydrolyze bacterial membrane phospholipids, which is a major structural component of the bacterial cell wall (27). Its relevance during infections stems from *in vivo* studies, in which mice overexpressing human hGIIA better control infections by group B Streptococcus, *Staphylococcus aureus* and *Bacillus anthracis* than their control littermates (41–44). Furthermore, the fact that pathogenic Gram-positive bacteria developed resistance against hGIIA is an indication for its relevance *in vivo* (44). Future work is needed to unravel whether serum sensitivity to hGIIA mediated killing is specific for commensal *E. faecium* strains or whether other non-pathogenic Gram-positives may be killed by normal human serum as well. Thus far, non-pathogenic Gram-positives appear resistant to serum killing (21). Only *B. anthracis* was previously shown to be sensitive to normal serum, however this was fully dependent on growth inhibition by transferrin-mediated iron deprivation (45). hGIIA is an acute phase reactant protein, whose serum concentration upon bacterial infection can increase up to hundred fold, which is high enough to kill some Gram-positive pathogens (31). Since hGIIA is able to kill commensal variants of a clinically relevant multi-resistant pathogen, and since hGIIA resistance can be an important mechanism of bacterial escape, strategies based on the use of components of the innate immune system such as hGIIA and even variants of this latter may be developed to fight against multidrug-resistant bacteria. In conclusion, the findings presented here not only provide fundamental knowledge about how the innate immune system kills bacterial cells, it also opens up new therapeutic routes to boost immune clearance of bacteria through conversion of serum resistance.

## Materials and Methods

### Bacterial strains and growth conditions

The 19 *E. faecium* strains used in this study are from our laboratory collection (department of medical microbiology, UMC Utrecht) and were previously characterized bacteria isolated from healthy and hospitalized humans or animals (Table 1) (9, 46). *E. faecium* was grown at 37°C for 24 hours in Trypticase soy agar II (TSA) plates supplemented with 5% sheep blood (BD Biosciences) and tryptic soy broth media (TSB; Oxoid) when indicated.

### Serum, plasma and inhibitors

Normal human serum (HS) was generated at the department of medical microbiology, UMC Utrecht. As previously described (21), whole blood was drawn via venous puncture from 20 healthy volunteers who provided written informed consent in accordance with the Declaration of Helsinki and approval was obtained from the medical ethics committee of the UMC Utrecht. Following collection via venous puncture, blood was clotted for 15 min at room temperature. Blood was centrifuged (10 min at 2700 × g at 4°C) and serum (supernatant) was collected, pooled and frozen in small aliquots before storage at −80°C. Heat-inactivated (HI) serum was prepared by incubating serum at 56°C for 30 min. When indicated, EDTA (10 mM), C3 inhibitor compstatin (CP40) (24) (10 µg/ml), C5 inhibitor OmCI (25) (10 µg/ml), group IIA secreted phospholipase A2 (hGIIA) inhibitor LY311727 (50 µM; Sigma-Aldrich Ldt.) were used. Pure recombinant hGIIA was prepared in *E. coli* as described (44, 47). hGIIA concentration in human serum was quantified using the human sPLA2-IIA enzyme linked immunosorbent assay (ELISA) kit from Cayman Chemicals (Ann Arbor, MI).

### Serum bactericidal assays

*E. faecium* strains were grown in 4 ml TSB to optical density 660 (OD660) 0.4 from an overnight culture in TSA plates supplemented with 5% sheep blood (BD Biosciences). Bacteria were centrifuged and resuspended in sterile RPMI (Gibco). For each specific condition, 10^5^ bacteria were incubated with human serum (at the indicated concentrations, with or without inhibitors) in sterile round-bottom 96-wells plates (Greiner) under shaking conditions for 30 min at 37°C. Killing was evaluated by serial dilution of samples in RPMI and subsequent plating onto TSA plates supplemented with 5% sheep blood. Following overnight incubation at 37°C, surviving bacteria were quantified by counting the number of colony forming units (CFU). Killing was measured by comparison of total colony forming units after 30 min incubation in the control (RPMI) in relation to 25% human serum or other described condition. Bactericidal assays were performed at least in duplicate per condition.

### Bacterial membrane permeabilization

Bacterial membrane permeability was analyzed by flow cytometry and confocal microscopy. Following incubation of *E. faecium* with serum, the membrane impermeable nuclei acid dye sytox green (Life technology, 1:5000 (v:v)) was added to the samples. Sytox green staining was quantified by flow cytometry on a BD FACSVerse (Becton Dickinson, San Jose, CA, USA, 488 nm laser). Bacteria were gated based on forward and side scatter properties and fluorescence of 10.000 bacterial cells was quantified. Results were analyzed with FlowJo (version v10). Based on the buffer negative control condition (RPMI), a threshold was set to determine the increase of sytox green signal in the population. The increase of sytox signal was represented as the percentage of the total population that was stained with sytox green (see results Fig 4A for example of flow cytometry plots). For confocal microscopy, bacteria were incubated with 25% serum as described above in the presence of syto9 (1.5 µl) and propidium iodide (1.5 µl) (both from BacLight bacteria viability kit, Life Technologies). After 10 min incubation at room temperature, bacteria were mounted in a glass slide with ProLong antifade mountant (Life Technologies), followed by fluorescence analysis in the confocal microscope (Leica SP5). Syto9 and propidium iodide were excited at 488 nm. Pictures were taken at 40x magnification and optical zoom of 3x.

### Surface deposition of activated complement products

Deposition of C3b molecules or C5b-9 complexes on *E. faecium* surface was quantified by flow cytometry as previously described (21). *E. faecium* was incubated with human serum (or plasma, with or without inhibitors) for 30 min at 37°C, shaking. After washing, bacteria were incubated with FITC-conjugated anti-C3 antibody (1 µg/ml, Protos Immunoresearch) or mouse anti-C5b-9 antibody (1 µg/ml, aE11 Santa Cruz) in PBS-1% BSA for 30 min at 4°C. For C5b-9 detection, a subsequent incubation with FITC-conjugated goat anti-mouse IgG antibody (1 µg/ml, Dako) was performed (30 min at 4°C). Bacteria were washed once more and fluorescence of 10.000-gated bacteria was quantified by flow cytometry using a FACSVerse flow cytometer (Becton Dickinson, San Jose, CA, USA) (48).

## Acknowledgements

We would like to thank Prof. Susan Lea from University of Oxford for kindly provide OmCI and Prof. John Lambris, University of Pennsylvania, Philadelphia for kindly providing Compstatin (CP40).

## Supplementary Material Captions

**S1 Figure. Complement system activity in human serum.** A. Killing of clinical *E. faecium* strain E1162 was measured by cell viability on blood agar plates after 30 min incubation on the indicated conditions. B. Inhibition of complement system was measured by C3b deposition on bacterial surface.

**S2 Figure. hGIIA concentration in pooled normal human serum.** hGIIA concentration in pooled normal human serum used in all assay was measured by Elisa kit (Cayman).

